# Individual differences in novelty-seeking behavior are modulated by GRK2/3 inhibition in rats

**DOI:** 10.64898/2026.09.25.753780

**Authors:** Simón Dávila, Catalina Starocelsky, Carolina Pino, Francisca Codocedo, Cristobal Guzmán-Muñoz, Benjamin Bossel, Francisco Novoa, Marcela González, María Estela Andrés, Rafael Ignacio Gatica

## Abstract

**Background and Purpose:** High novelty-seeking is an endophenotype associated with greater vulnerability to drug addiction, marked by a hyperdopaminergic state and reduced dopamine D2 receptor (D2R) levels in the dorsal striatum (DS). G protein-coupled receptor kinases (GRKs) regulate desensitization, internalization, and recycling of these receptors, including the D2R. This study investigated the relevance of GRK2 and 3 (GRK2/3) in novelty-seeking behavior and striatal dopamine dynamics.

**Experimental Approach:** Individual novelty-seeking trait was assessed in Sprague Dawley rats using an open-field test. D2R, GRK2, and GRK5 protein levels were measured in synaptosomal DS samples using Western Blot. Pharmacological inhibition of GRK2/3 was assessed using Cmpd101. We tested the effect of systemic Cmpd101 administration on contextual novelty discrimination and local Cmpd101 perfusion on baseline and stimulus-evoked extracellular dopamine (DA) levels in the DS.

**Key Results:** Locomotor activity in a novel environment inversely correlated with the synaptosomal GRK2 and D2R protein levels in the DS. Systemic Cmpd101 treatment modulated the locomotor discrimination index in a novelty phenotype-dependent manner. Moreover, local Cmpd101 perfusion in the DS elicited a sustained elevation in basal extracellular dopamine levels and significantly blunted subsequent stimulus-evoked DA release, both effects independent of novelty-seeking phenotype.

**Conclusion and Implications:** Together, our findings link GRK2/3 signaling to individual differences in novelty-seeking and striatal dopaminergic function. These results suggest a potential relevance to drug addiction vulnerability.

## Introduction

Drug addiction is a chronic relapsing disorder characterized by compulsive drug seeking and consumption despite adverse consequences (Koob & Volkow, 2016). Although many individuals are exposed to addictive substances, only a subset develops substance use disorders, pointing to individual vulnerability factors shaped by genetic, molecular, and neurobiological differences (Egervari et al., 2018). Importantly, only ∼15-17% of individuals exposed to addictive drugs progress toward compulsive and chronic drug use, a phenomenon that is similarly reproduced in rodent models of addiction vulnerability (Anthony et al., 1994; Deroche-Gamonet et al., 2004). Novelty-seeking behavior, defined as the tendency to desire and explore novel stimuli, is one of the most robust behavioral endophenotypes predicting the acquisition of drug consumption, the transition to compulsive use, and a higher propensity to relapse (Belin & Deroche-Gamonet, 2012; Wingo et al., 2016). To investigate the neurobiological underpinnings of this phenomenon, the High Responder (HR) and Low Responder (LR) rat paradigm effectively captures this individual vulnerability based on locomotor activity to a novel environment (Piazza et al., 1989). While this novelty-seeking trait robustly predicts enhanced susceptibility to drug-induced plasticity and addiction-like behaviors (Flagel et al., 2014), the underlying synaptic mechanisms driving these divergent phenotypes remain an open question. The transition to compulsive behaviors is accompanied by a functional reorganization of dopaminergic control within the dorsal striatum (DS) (Everitt & Robbins, 2016). Consequently, maintaining the precise regulation of dopaminergic signaling specifically within the DS is pivotal to prevent the aberrant reinforcement of found in drug addiction.

To maintain optimal circuit function, this striatal dopaminergic tone is tightly autoregulated by dopamine D2 receptors (D2Rs). While postsynaptic D2Rs on medium spiny neurons mediate motor output, a distinct population of presynaptic D2 autoreceptors provides an essential inhibitory feedback loop (Ford, 2014). Activation of these autoreceptors strictly suppresses further dopamine (DA) synthesis and release, functioning as a critical homeostatic brake (Franco et al., 2021). Notably, individual differences in novelty-seeking are associated with distinct neurochemical profiles, including variations in basal DA tone (Mabrouk et al., 2018).For instance, LR rats exhibit higher striatal D2R density compared to their highly reactive HR counterparts (Hooks et al., 1994).

The functional availability and signaling of these D2Rs are dynamically regulated by homologous desensitization mediated by G protein-coupled receptor kinases (GRKs), specifically the β-adrenergic receptor kinases GRK2 and GRK3 (for now on, GRK2/3) (Gurevich et al., 2012). Upon DA binding, GRK2/3 phosphorylate the third intracellular loop of the D2R, promoting β-arrestin recruitment and subsequent receptor internalization (Beaulieu & Gainetdinov, 2011). In vivo, genetic models demonstrate that distinct GRK isoforms exert highly specific, cell-type and region-dependent effects on DA neurotransmission (Gurevich et al., 2016). For instance, GRK2 deletion in D1R or D2R-expressing neurons produces opposing effects, i.e., an increase or a decrease in stimulated DA levels in the striatum, respectively (Daigle et al., 2014). Moreover, GRK3 deficiency increases DA release induced by amphetamine (Sellgren et al., 2021). However, whether GRK-mediated regulation directly dictates trait novelty-seeking behavior and associated striatal DA levels remains unknown.

In the present study, we hypothesized that individual differences in novelty-seeking are associated with the synaptic abundance of GRK2/3 and D2R in the DS, and that pharmacological inhibition of these kinases alters both novelty-seeking behavior and DA dynamics. To test this, we first investigated the relationship between locomotor activity in a novel environment and the synaptic protein levels of D2R, GRK2, and, as a specificity control, the GRK5 isoform. Subsequently, we utilized Cmpd101, a highly selective and brain-penetrant inhibitor of GRK2/3 (Thal et al., 2011), to evaluate how acute systemic blockade of these enzimes modulates contextual novelty discrimination. Finally, employing *in vivo* microdialysis, we determined how local GRK2/3 inhibition in the DS alters absolute and stimulus-evoked DA release, and whether the efficacy of this neurochemical regulation is dictated by the intrinsic novelty-seeking phenotype.

## Materials and Methods

### Animals

Adult male Sprague-Dawley rats (9 weeks old, 300-500 g at the start of the experiments) were used in this study and obtained from the CIBEM animal facility (Center for Innovation in Biomedical Experimental Models) at the Pontificia Universidad Católica de Chile. Animals were housed in pairs under controlled environmental conditions, with a 12-h light/dark cycle (lights on at 09:00 am), constant temperature (22–24 °C), and *ad libitum* access to food and water. Upon arrival, rats were transferred to the Neuroepigenetics Laboratory facilities and allowed a 1-week habituation and handling period prior to the start of experimental procedures. All experimental protocols were reviewed and approved by the Institutional Animal Care and Use Committee (CEC-CAA) of the Pontificia Universidad Católica de Chile (protocol code: 220420014) and were conducted in accordance with the “NIH Guide for the Care and Use of Laboratory Animals” (8th Edition) and the “Guidelines for the Use of Animals in Neuroscience Research” of the Society for Neuroscience.

### Drugs and Reagents

The selective GRK2/3 inhibitor Cpmd101 (Cmpd; Hello Bio, United Kingdom) was used for both systemic behavioral testing and local *in vivo* microdialysis. For systemic intraperitoneal (i.p.) administration, Cmpd101 was dissolved at a concentration of 1.0 mg/mL in a vehicle solution consisting of 90% saline (0.9% NaCl; Laboratorio Sanderson S.A., Chile), 5% ethanol (95% ethanol; Winkler Ltda., Chile), and 5% Kolliphor (Sigma-Aldrich, Germany), and administered at a dose of 1.0 mg/kg. Saline (0.9% NaCl) was used for injection adaptation days. For local intra-striatal reverse dialysis, Cmpd101 was prepared from a 10 mM stock solution (dissolved in 100% DMSO) and diluted to a final concentration of 100 µM in Krebs-Ringer phosphate (KRP) buffer.

### Open field test

Following the 1-week habituation and handling period, individual differences in novelty-seeking were assessed in an open-field arena. Each session lasted 1 h and was conducted in a black arena (100 × 100 cm), with behavior recorded using an overhead camera (Logitech C270 HD). Locomotor activity was then analyzed using ANY-maze software (Stoelting Co., USA). Testing was performed between 9:00 and 15:00 h under standard room lighting conditions. Twenty four hours after behavioral testing, rats were euthanized and brains were extracted.

For the novelty discrimination paradigm, locomotor activity under familiar versus novel conditions was assessed using a modified open-field apparatus with interchangeable floor inserts. The protocol spanned 11 days. During the first 5 consecutive days, animals were placed in the arena with a uniform black floor (familiar context) to establish baseline locomotion and promote habituation. On days 8 and 9, animals received i.p. saline injections to account for potential handling and injection stress. Pharmacological testing was conducted on days 10 and 11. On the first test day, animals received either vehicle or Cmpd101 (1.0 mg/kg, i.p.), followed by a 30-min interval before being placed in the habituated open field (Context 1) for 60 min. On the second test day, animals underwent the identical injection procedure but were exposed to a visually novel environment featuring a patterned floor with two 50 x 50 cm cyan squares positioned in opposite corners (novel context). To prevent order effects, the presentation of Context 1 and Context 2 during the testing phase was counterbalanced across subjects. Locomotor activity in black and cyan zones, and velocity were analyzed with ANY-maze software (Stoelting Co., USA). The locomotor discrimination index (LDI) was used as a parameter for novelty exploration as follows: LDI = (Locomotor activity in cyan − black) / total locomotor activity. Two hours after the last treatment injection, rats were euthanized and brains were extracted.

### Synaptosomal Protein Extraction and Western Blotting

DS and nucleus accumbens (NAc) were dissected and synaptosomal and cytosolic fractions were isolated using Syn-PER™ Synaptic Protein Extraction Reagent (Thermo Scientific, USA) supplemented with protease and phosphatase inhibitors. Tissue was homogenized and centrifuged at 1,200 x g for 10 min at 4 °C. The supernatant was subjected to a second centrifugation at 15,000 x g for 20 min at 4 °C to yield the synaptosomal pellet, which was resuspended and stored at −80 °C. Protein concentrations were determined using a micro-BCA Protein Assay Kit (Thermo Scientific, USA).

Proteins (20 µg per sample) were denatured at 100 °C for 10 min in sample buffer containing SDS 2%, glycerol 10%, bromophenol blue 0.0025%, and β-mercaptoethanol 5%, separated on 8% SDS-PAGE gels, and transferred onto 0.2 µm nitrocellulose membranes (Bio-Rad Laboratories, USA) at 400 mA for 2 h at 4 °C. Membranes were stained with Ponceau S solution (0.2% in 5% acetic acid) to quantify total protein content for normalization, imaged using a GeneGnome XRQ system (Syngene, United Kingdom), and sectioned by molecular weight. After blocking in 5% non-fat dry milk in TBS-Tween 0.1% for 1 h, membranes were incubated overnight at 4 °C with primary antibodies: mouse anti-GRK2 (1:5000; sc-13143, Santa Cruz Biotechnology, USA), mouse anti-GRK5 (1:5000; sc-518005, Santa Cruz Biotechnology, USA), and mouse anti-D2R (1:1000; sc-13143, Santa Cruz Biotechnology, USA). Membranes were subsequently incubated with HRP-conjugated donkey anti-mouse secondary antibody (1:5000; 715-035-150, Jackson ImmunoResearch, USA) for 1 h at room temperature. Immunoreactive bands were detected using SuperSignal West Pico (Thermo Fisher, USA) and the GeneGnome system. Band intensities for GRK2, GRK5, and D2R were quantified using ImageJ software.

### Stereotaxic Surgery and In Vivo Microdialysis

On the day following the initial open-field phenotyping, animals underwent stereotaxic surgery for microdialysis. Anesthesia was induced with urethane (1.5 g/kg, i.p.; Sigma-Aldrich, Germany) administered in two divided doses to ensure a stable surgical plane. Animals were secured in a stereotaxic frame (Stoelting Co., USA) and the skull was exposed and leveled. A craniotomy and subsequent durectomy were performed to unilaterally lower a microdialysis probe (Cat. n° MAB 2.14.2, Microbiotech, Sweden) into the DS. Coordinates were determined based on the Paxinos & Watson (2009) rat brain atlas: AP +1.2 mm, ML +3.6 mm, and DV −5.0 mm under dura.

The probe was connected to a perfusion pump (Harvard Instruments, USA). The standard perfusate consisted of a basal Krebs-Ringer phosphate buffer (KRP-N; pH 7.4) composed of 120 mM NaCl, 2.4 mM KCl, 1.2 mM CaCl_₂_, 0.9 mM NaH_₂_PO_₄_, and 1.4 mM Na_₂_HPO_₄_, supplemented with 1% DMSO. For depolarizing stimulations, a high-potassium buffer (KRP-K) was utilized, containing 40 mM KCl with NaCl adjusted to 82.4 mM to maintain osmolarity.

KRP perfusion was carried out at a constant flow rate of 2 µL/min. Following probe insertion, a 40-min stabilization period was allowed. Dialysates were collected in 10-min fractions (20 µL per fraction) into tubes containing 4 µL of 0.2 N perchloric acid (PCA; Merck, Germany) and kept on ice. After stabilization, five baseline dialyzate samples were collected during KRP-N perfusion. Subsequently, group-specific protocols were applied:

- **Vehicle Group:** A first depolarizing stimulation (S1) with KRP-K was applied (10 min), followed by KRP-N (50 min), a second stimulation (S2) with KRP-K (10 min), and a final reperfusion with KRP-N (30 min).
- **Cmpd101 Group:** A first depolarizing stimulation (S1) with KRP-K was applied (10 min), followed by KRP-N (20 min). Next, KRP-N supplemented with 100 µM Cmpd101 was perfused (30 min), followed by a second stimulation (S2) with KRP-K + Cmpd101 (10 min), and a final reperfusion with KRP-N + Cmpd101 (30 min).

Upon experiment completion, brains were extracted and fixed in 4% paraformaldehyde for histological verification of probe placement (Supplementary Figure 1).

### Dopamine Quantification

DA levels in dialysate samples were quantified using high-performance liquid chromatography with electrochemical detection (HPLC-ECD). A 20 µL volume from each sample was manually injected (Cat. n° 7725i, Rheodyne). The system comprised a dual-piston pump (PU-2089S Plus, Jasco), a silica gel column (5020-04424, GL Sciences), and an amperometric detector (LC-4C, BASi) with the working electrode oxidation potential set to +650 mV. The mobile phase consisted of 96.96 mM NaH_₂_PO_₄_, 1 mM 1-octanesulfonic acid, 1 mM EDTA, and 5.5% (v/v) acetonitrile, adjusted to pH 4.0. The system was operated under isocratic conditions at a flow rate of 0.5 mL/min, yielding a DA retention time of 12-14 min.

### Statistical Analysis

All data processing, statistical analyses and figure generation were conducted using Python (version 3.12.3), employing the SciPy, statsmodels, and Pingouin libraries.

Western blot analyses were performed using linear mixed models, as in Omondi et al., 2024. Analyses were conducted separately for each protein target and, for the locomotor analysis, each brain region and subcellular fraction. Lanes with nonpositive protein signal or Ponceau signal ≤0.05 were excluded. Within each animal and target, lanes with an absolute modified median absolute deviation score >2.5 for the protein-to-Ponceau ratio were also excluded. All remaining lanes entered the models. Protein and Ponceau signals were scaled by their respective target-specific means for numerical stability. Ponceau signal was included as a covariate; the protein-to-Ponceau ratio was used only to identify outlying lanes. To assess the association between day 1 locomotor activity and protein abundance, the model included locomotor activity and Ponceau signal as fixed effects and crossed random intercepts for animal and Western blot membrane. Locomotor activity was centered on the mean across animals. The locomotor slope was evaluated with a two-sided Wald z test. To compare GRK2 protein levels between Cmpd101 with vehicle treatments, model included treatment and Ponceau signal as fixed effects, with animal and membrane random intercepts where estimable. Vehicle was the reference group. The adjusted treatment effect was evaluated with a one-degree-of-freedom Wald χ² test. Model estimates and 95% confidence intervals were reported.

Behavioral analyses compared Cmpd101 with vehicle groups. For each outcome, the groups were compared using a two-sided Welch’s t test. Separate ordinary least-squares (OLS) models included treatment, day 1 baseline locomotor activity, and their interaction. Baseline activity was centered on the mean across analyzed animals. These models used heteroskedasticity-consistent (HC3) robust standard errors and a finite-sample t reference. Group differences and model coefficients were reported with 95% confidence intervals.

Absolute DA concentrations at 60 min, before vehicle or Cmpd101 perfusion, and 90 min, after perfusion, were analyzed using a mixed analysis of variance with time as the within-animal factor and treatment as the between-animal factor. Comparisons between treatments at each time point and for the S2/S1 ratio used Welch’s t tests. Changes from 60 to 90 min within each treatment group were assessed with paired t tests. The association between baseline dopamine concentration and day 1 locomotor activity was assessed using Pearson correlation. Separate ordinary least-squares models assessed dopamine at 60 min, dopamine at 90 min, and the S2/S1 ratio as functions of locomotor activity, treatment, and their interaction. Locomotor activity was mean-centered. Models used HC3 robust standard errors and a finite-sample t reference.

## Results

### Synaptic GRK2 and D2R protein levels in the DS inversely associates with novelty-seeking behavior

We investigated whether individual differences in the behavioral response to a novel environment associate with the synaptic levels of these proteins. We characterized the locomotor phenotype of a large cohort of Sprague-Dawley rats (n = 50) and subsequently isolated dorsal striatal (DS) synaptosomes to quantify GRK2, GRK5, and D2 receptor (D2R) protein levels (Fig. 1A). Behavioral characterization revealed a broad continuous distribution of locomotor scores, reflecting substantial inter-individual variability in novelty-induced locomotor activity (Fig. 1B). Representative Western blot analysis illustrated the baseline protein levels within isolated synaptosomes, capturing the spectrum of subjects with low versus high locomotor activity (Fig. 1C). Biochemical quantification revealed a strong inverse relationship between locomotor activity and synaptic D2R abundance, where high locomotor activity was highly predictive of reduced synaptosomal D2R protein levels (Fig. 1D; p < 0.001, LMM). Interestingly, a significant inverse relationship was also observed for GRK2; specifically, greater distances traveled in the novel environment robustly predicted lower levels of synaptosomal GRK2 (Fig. 1E; p < 0.001, LMM). In contrast, synaptosomal levels of GRK5 showed no association with individual novelty seeking (Fig. 1F; p = 0.236, LMM). Together, these data demonstrate that individual differences in novelty seeking are associated with synaptic abundance of GRK2 and D2R in the DS.

**FIGURE 1.**
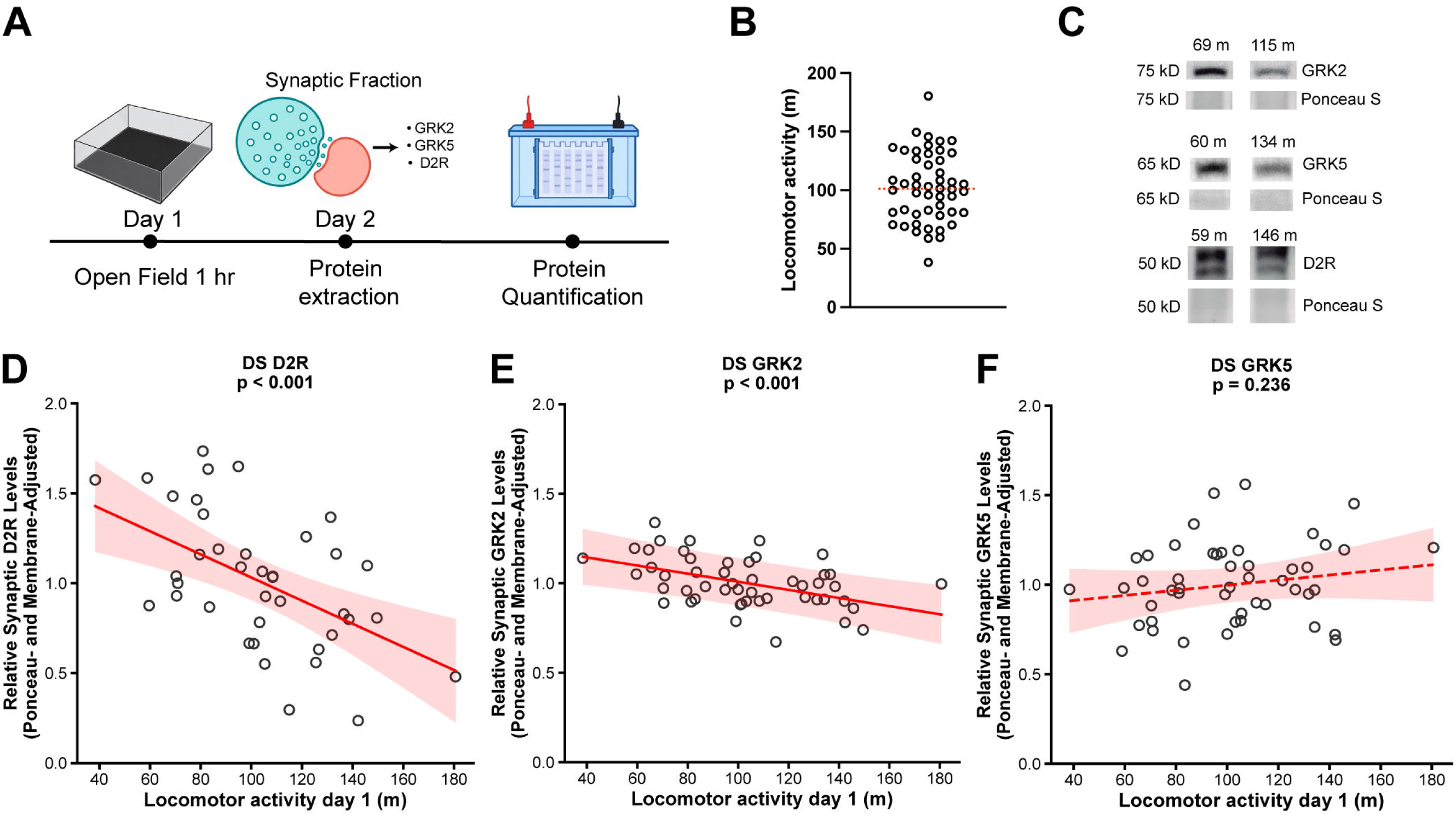
Synaptic D2R and GRK2 protein levels in the DS inversely associate with novelty-seeking behavior. A) Experimental timeline for behavioral phenotyping and biochemical analysis. On Day 1, rats were individually placed in an open-field arena (100 × 100 cm) for 1 hour; total distance traveled was recorded as the locomotor response to novelty. The next day, brain regions were dissected and subjected to subcellular fractionation to obtain synaptosomal and cytosolic fractions. GRK2, GRK5, and D2R protein levels were subsequently quantified by Western blot. B) Population distribution of locomotor activity on Day 1 (n = 50). The dashed red line indicates the population median (101.2 m). C) Representative Western blots in the dorsal striatal synaptosomal fraction. Western blot images of GRK2 (75 kDa), GRK5 (65 kDa), and D2R (50 kDa) are shown for pairs of animals with contrasting locomotor activity levels (GRK2: 69 m vs. 115 m; GRK5: 60 m vs. 134 m; D2R: 59 m vs. 146 m). Ponceau S staining is shown as the corresponding total protein loading control. D) Synaptosomal D2R levels inversely associate with individual locomotor activity (n = 39; p < 0.001). E) Synaptosomal GRK2 levels inversely associate with individual locomotor activity (n = 50; p < 0.001). F) Synaptosomal GRK5 levels do not associate with individual locomotor activity (n = 49; p = 0.236). Linear mixed models (LMMs) were utilized to account for inter-membrane variability, with Ponceau S signal included as a covariate to control for total protein loading. Solid red lines (D-E) denote statistically significant correlations, whereas the dashed red line (F) indicates a non-significant relationship. Shaded areas represent 95% confidence intervals, and each open circle represents one individual animal.

To determine the anatomical and subcellular specificity of these findings, we also measured protein levels in the corresponding DS cytosolic fraction, as well as in the synaptosomal fraction of the nucleus accumbens (NAc). Biochemical quantification revealed no significant associations in these additional compartments. Specifically, novelty-induced locomotion showed no relationship with cytosolic DS levels of GRK2 (Supplementary Fig. 2A; p = 0.542, LMM), GRK5 (Supplementary Fig. 2B; p = 0.349, LMM), or D2R (Supplementary Fig. 2C; p = 0.976, LMM). Furthermore, synaptic protein levels in the NAc exhibited no association with novelty seeking for either GRK2 (Supplementary Fig. 2D; p = 0.658, LMM) or GRK5 (Supplementary Fig. 2E; p = 0.59, LMM). Together, these data demonstrate that the inverse relationship between individual novelty-seeking behavior and GRK2/D2R levels is highly compartmentalized, being specifically restricted to the synapses of the DS.

### Systemic GRK2/3 inhibition effects on contextual novelty discrimination are novelty-seeking dependent

To evaluate the behavioral relevance of these neurochemical associations, we investigated whether pharmacological inhibition of GRK2/3 alters novelty seeking. The experimental design comprised initial habituation to a familiar environment (Days 1–5), followed by procedural acclimation via intraperitoneal (i.p.) saline injections (Days 6–7), and treatment testing on Days 8 and 9. On testing days, subjects received an i.p. injection of either Vehicle or the GRK2/3 inhibitor Cmpd101 (1.0 mg/kg) prior to context exposure, with the presentation of familiar and novel contexts counterbalanced across both days (Fig. 2A). Prior to context testing, animals exhibited a continuous distribution of baseline locomotor activity on Day 1 (Fig. 2B). To robustly quantify this response, we calculated the Locomotor Discrimination Index (LDI), a metric designed to evaluate an animal’s capacity for contextual novelty discrimination by measuring the preference for exploring a novel environment relative to a familiar one, normalized by total activity. At the cohort level, Cmpd101 administration did not alter the overall locomotor activity in the familiar or novel context (Fig. 2C-D; p = 0.500 and p = 0.978, respectively, Welch’s independent-samples t-test), and the LDI remained unchanged (Fig. 2E; p = 0.248, Welch’s independent-samples t-test). However, incorporating locomotor activity on day 1 into the analysis via OLS + HC3 regression revealed a distinct behavioral profile. Although GRK2/3 inhibition did not significantly affect absolute locomotor activity in either context as a function of the intrinsic phenotype (Fig. 2F-G; main treatments and interactions p > 0.05, OLS regression + HC3), it significantly modified the relationship between baseline novelty seeking and contextual exploration. Specifically, a significant Treatment × Locomotor Activity interaction was found for the LDI (Fig. 2H; OLS regression + HC3; interaction p = 0.030), with no overall treatment effect (treatment effect p = 0.260, OLS regression + HC3). This means that animals with a high novelty-seeking increase their novelty discrimination in the novel open field with GRK2/3 inhibition. Taken together, these data demonstrate that the behavioral efficacy of GRK2/3 inhibition is strictly dictated by the subject’s intrinsic locomotor phenotype, functioning as a critical, phenotype-specific modulator of contextual novelty discrimination.

**FIGURE 2.**
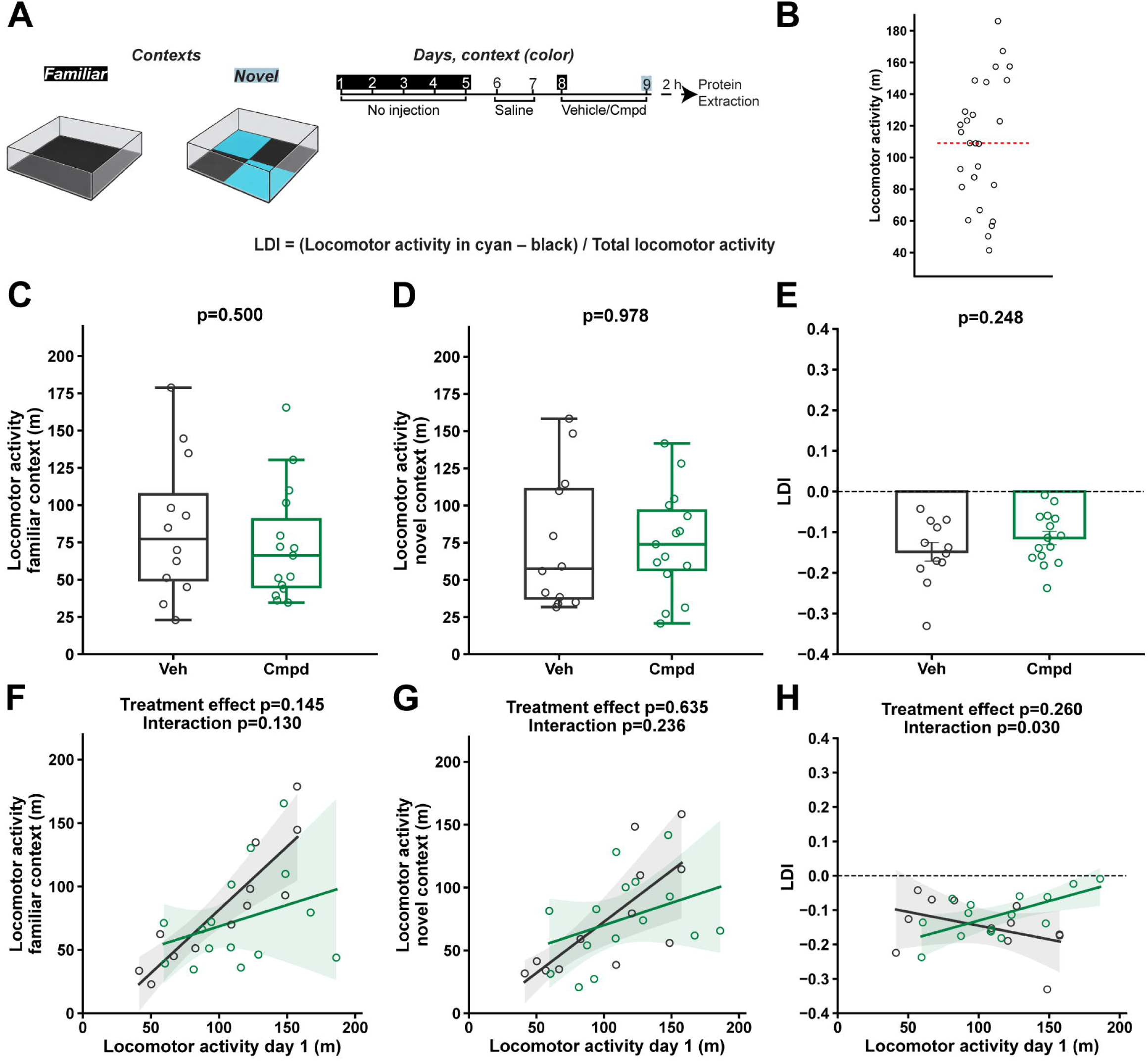
Cmpd101 effects on contextual locomotor discrimination depend on baseline novelty-seeking. A) Experimental protocol for contextual discrimination. Rats were exposed to an open field with a black floor over five consecutive days (familiar context, noted by black shading in the protocol day numbers). On days 6 and 7, rats were injected with saline (i.p.) and returned to their home cages. On days 8 and 9, rats were injected with Vehicle or Cmpd101 (1.0 mg/kg, i.p.), and their behavior was tested in either a familiar or novel context (cyan floor). Context presentation order was counterbalanced across animals. Locomotor Discrimination Index (LDI) was calculated as: (Locomotor activity in cyan − black) / total locomotor activity. B) Population distribution of locomotor activity on Day 1. The dashed red line indicates the median. C) Cmpd101 does not affect locomotor activity in the familiar context (p = 0.500). D) Cmpd101 does not affect locomotor activity in the novel context (p = 0.978). E) Locomotor discrimination index is maintained across treatment groups (p = 0.248). Group differences were analyzed using Welch’s independent-samples t-test. Box-and-whisker plots represent the median and interquartile range. F) Cmpd101 did not modulate locomotor activity in the familiar context as a function of locomotor activity on day 1 (Treatment p = 0.145; Interaction p = 0.130). G) Cmpd101 did not modulate locomotor activity in the novel context as a function of locomotor activity on day 1 (Treatment p = 0.635; Interaction p = 0.236). H) Cmpd101 differentially modulates LDI as a function of locomotor activity on day 1 (Treatment p = 0.260; Interaction p = 0.030). Baseline-dependent analyses utilized ordinary least-squares (OLS) regressions with heteroskedasticity-consistent standard errors (HC3). Black and green lines represent Vehicle and Cmpd101 groups, respectively, and shaded areas represent 95% confidence intervals of the mean prediction. The dashed horizontal line at y = 0 demarks the threshold of no discrimination between contexts. Individual data points are shown across all relevant panels (Vehicle: n = 12; Cmpd101: n = 15).

To rule out the possibility that the phenotype-dependent changes in contextual discrimination were secondary to motor impairments, we evaluated active movement speed across both testing environments. At the cohort level, Cmpd101 administration did not alter active movement speed in either the familiar (Supplementary Fig. 3A; p = 0.277, Welch’s independent-samples t-test) or the novel context (Supplementary Fig. 3B; p = 0.303, Welch’s independent-samples t-test). Furthermore, incorporating locomotor activity on day 1 into the analysis revealed that movement dynamics remained completely unaffected. Although GRK2/3 inhibition modulated the LDI, active movement speed in both the familiar (Supplementary Fig. 3C; Treatment effect p = 0.452, Interaction p = 0.683, OLS regression + HC3) and novel contexts (Supplementary Fig. 3D; Treatment effect p = 0.409, Interaction p = 0.850, OLS regression + HC3) was entirely independent of both Cmpd101 treatment and the intrinsic locomotor phenotype.

To determine whether systemic GRK2/3 inhibition modifies GRK2 levels in the brain, DS samples were collected 2 hours after treatment injection and processed for Western Blot (Supplementary Figure 4). Data showed a decrease in GRK2 protein levels in the Cmpd group compared to the vehicle group (p = 0.006, Wald chi-square test). This result indicates that GRK2/3 inhibition reduces GRK2 availability in the DS, which might relate to the changes found in contextual novelty discrimination.

### GRK2/3 inhibition in the DS alters DA dynamics independently of novelty-seeking behavior

To determine whether GRK2/3 inhibition alters striatal function in these rats with differential novelty seeking behavior, we performed *in vivo* microdialysis in the dorsolateral striatum. The experimental timeline covered an initial Open Field assessment (Day 1), followed by microdialysis probe and DA measurement on anesthetized rats (Day 2) (Fig. 3A). Following a stabilization period, the *in vivo* sampling protocol was executed, consisting of baseline sampling, a first depolarizing stimulus (S1) with K^+^ 40 mM, intra-microdialysis probe local perfusion of the assigned treatment (Vehicle or Cmpd101), and a second depolarizing stimulus (S2) under treatment (Fig. 3B). Analysis of DA release dynamics via representative time-course profiles revealed a marked attenuation of the peak response to S2 specifically in Cmpd101-treated subjects (Fig. 3C).

**FIGURE 3.**
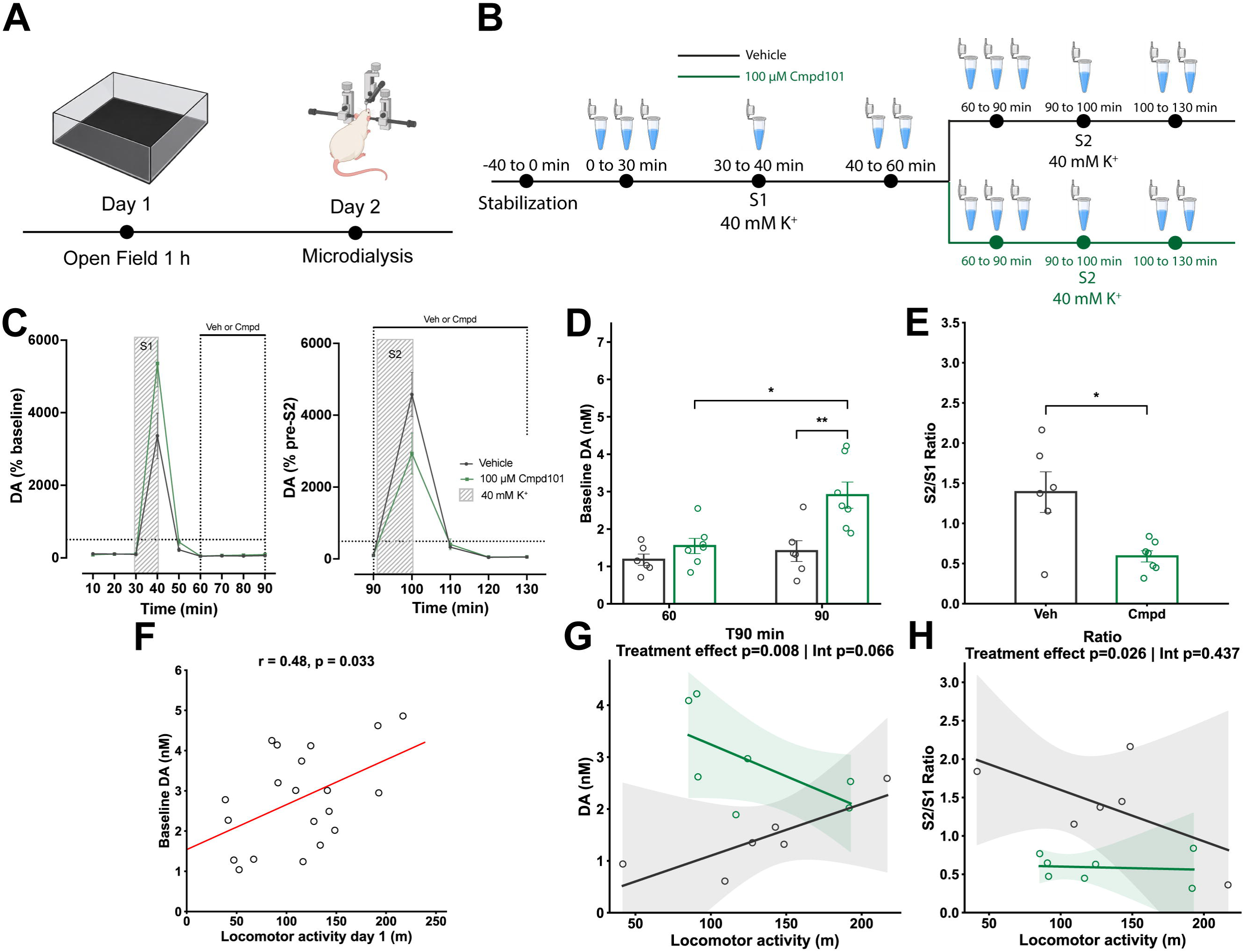
Baseline dorsolateral striatal dopamine correlates with novelty-seeking behavior, whereas Cmpd101 alters dopamine dynamics independently of this intrinsic phenotype. **A)** Experimental timeline for microdialysis and behavioral testing. Day 1: Locomotor activity testing in an open-field arena (100 × 100 cm). Day 2: Microdialysis in the DS. B) Microdialysis session protocol and pharmacological treatment perfusion. The baseline perfusion buffer consists of Krebs-Ringer phosphate buffer with normal potassium (2.4 mM, KRP-N) containing 1% DMSO. The depolarizing stimulations (S1 and S2) consist of KRP with 40 mM potassium (KRP-K) containing 1% DMSO. Sequential dialysate collection occurred during an initial baseline (0-30 min), a first depolarizing pulse with KRP + 40 mM K^+^ (S1, 30-40 min), and an intermediate recovery period (40-60 min). During local perfusion of Vehicle (black line) or Cmpd101 (green line), a subsequent baseline (60-90 min), a second depolarizing pulse with KRP + 40 mM K^+^ (S2, 90-100 min), and a final recovery period (100-130 min) were collected. **C)** Time course of stimulus-evoked DA efflux relative to baseline (mean DA levels between 0-30 min). Extracellular DA during the first (S1, left) and second (S2, right) stimulation pulses are expressed as a percentage of their respective pre-stimulation baselines (60 min sample for S2). Hatched boxes indicate stimulation duration. Data are plotted as mean ± SEM for illustrative purposes. **D)** Cmpd101 increases extracellular DA levels at T90 compared to Vehicle (p = 0.007) and pre-treatment baseline T60 (p = 0.022). **E)** Cmpd101 diminishes the capacity for stimulus-evoked DA release (p = 0.024). Statistical significance was determined by a mixed two-way ANOVA followed by prespecified Welch’s independent and paired t-tests (panel D), and Welch’s independent-samples t-test (panel E). Bars represent mean ± SEM, and individual data points are shown (Vehicle = 6 rats, Cmpd101 = 7 rats). Significant differences are indicated by asterisks (*p < 0.05, p < 0.01). **F)** Baseline dorsolateral striatal DA levels positively correlate with novelty-induced locomotor activity (r = 0.48; p = 0.033, Pearson correlation). **G)** Cmpd101 effect on DA levels was independent of locomotor activity on Day 1 (Treatment p = 0.008; Interaction p = 0.066). **H)** Cmpd101 effect on evoked DA release was independent of locomotor activity on Day 1 (Treatment p = 0.026; Interaction p = 0.437). Ordinary least-squares (OLS) regressions with heteroskedasticity-consistent standard errors (HC3) was used in panels G-H. The solid red line represents the linear correlation fit. Black and green lines represent Vehicle and Cmpd101 groups, respectively, and shaded areas represent 95% confidence intervals of the mean prediction. Open circles represent individual animals.

To quantify this effect, we first evaluated extracellular DA levels during local treatment perfusion. Cmpd101 significantly increased absolute DA concentrations following 30 minutes of perfusion (T90) compared to the Vehicle group (Fig. 3D, p = 0.007), as well as compared to its own pre-treatment baseline at T60 (Fig. 3D, p = 0.022, mixed two-way ANOVA followed by prespecified pairwise t-tests). Importantly, local perfusion of Cmpd101 without a previous stimulation did not modify DA extracellular levels (Supplementary Fig. 5; p = 0.8366), indicating that the drug-induced elevation observed at T90 is contingent upon prior depolarization of the DA system. Subsequently, we assessed stimulus-evoked release by calculating the S2/S1 ratio, defined as the relative percentage change of the DA levels during the S2 stimulus divided by the S1 response. This analysis demonstrated a significant suppression of evoked release in animals subjected to GRK2/3 inhibition (Fig. 3E; p = 0.024, Welch’s independent-samples t-test).

Integrating these neurochemical alterations with the novelty seeking trait, we found that baseline dopaminergic tone in the DS is positively correlated with Day 1 locomotor activity (Fig. 3F, r = 0.48, p = 0.033, Pearson correlation). However, regression analysis revealed that the neurochemical impact of Cmpd101 treatment occurs independently of locomotor activity on Day 1. Both the increase in absolute DA at T90 (Fig. 3G; Treatment effect, p = 0.008, Interaction p = 0.066, OLS regression + HC3) and the decrease of evoked release (S2/S1 ratio) (Fig. 3H, Treatment effect, p = 0.026, Interaction p = 0.437, OLS regression + HC3) did not show an interaction with locomotor activity on Day 1. These findings suggest that while GRK2/3 inhibition exerts a generalized suppressive effect on dopaminergic dynamics, this effect is independent of novelty seeking phenotypes.

## Discussion

The present study identifies synaptic GRK2 in the DS as a critical molecular correlate of individual differences in novelty-seeking behavior and demonstrates that acute GRK2/3 inhibition alters dopaminergic dynamics in the DS while differentially affecting contextual novelty discrimination based on pre-existing behavioral phenotypes. Together, these findings identify the GRK2/D2R regulatory cascade in the DS as a relevant mechanism modulating both intrinsic novelty activity and striatal neurochemical dynamics.

Our biochemical data extends previous evidence linking novelty-seeking to differences in striatal dopaminergic function. The lower synaptic D2R abundance observed in animals with high novelty-induced locomotion aligns with prior studies reporting reduced striatal D2R density in individuals highly responsive to novelty (Hooks et al., 1994; Tournier et al., 2013). More broadly, this observation is consistent with preclinical and clinical evidence consistently associating reduced D2R availability with phenotypes that confer vulnerability to drug addiction (Dalley et al., 2007; Nader et al., 2006; Volkow et al., 1993). Importantly, discovering an inverse association for GRK2, while GRK5 showed no relationship, highlights an isoform-specific involvement in this behavioral trait. Given the established role of GRK2/3 in D2R homologous desensitization and trafficking (Beaulieu & Gainetdinov, 2011; Namkung et al., 2009), these associations suggest that behavioral vulnerability is accompanied not only by a reduction in receptor abundance but also by a constitutive deficit in the enzymatic machinery required to regulate dopaminergic function. The anatomical and subcellular specificity of these associations further underscores their functional relevance. The molecular correlations with novelty-seeking were detected exclusively in the synaptosomal fraction of the DS, with no significant associations observed in the corresponding cytosolic fraction or within the NAc. This regional specificity is particularly relevant, as the DS is progressively recruited during the transition from goal-directed drug seeking to habitual and compulsive behavior patterns (Everitt & Robbins, 2016).

Systemic administration of the selective GRK2/3 inhibitor Cmpd101 (Thal et al., 2011) did not alter overall locomotor activity or active movement speed at the population level, ruling out nonspecific motor or sedative confounds. However, its behavioral impact was critically dictated by the animal’s baseline novelty seeking. Regression analyses revealed that GRK2/3 inhibition enhanced novelty discrimination predominantly in animals exhibiting a high novelty-seeking phenotype. Because these highly novelty-driven animals show lower GRK2 and D2R protein levels at the synaptic levels, their dopaminergic circuits may operate near a threshold of dysregulation. Consequently, acute pharmacological inhibition of their GRK2/3 machinery alters how these distinct behavioral endophenotypes and respond to contextual novelty.

To further examine the role of GRK2/3 in striatal dopaminergic function, local GRK2/3 inhibition in the DS revealed a disruption of extracellular DA dynamics during repeated depolarizing stimulations. Following an initial depolarizing stimulus, intra-DS perfusion of Cmpd101 induced a sustained increase in basal extracellular DA levels compared to the vehicle group. Moreover, the dopaminergic response to a second depolarizing stimulus was significantly attenuated. Since extracellular DA levels reflect the balance between DA release and clearance, the present experiments do not establish which of these specific processes’ accounts for the observed effects. Nevertheless, the sustained elevation in extracellular DA followed by an attenuated response to subsequent stimulation suggests that GRK2/3 contributes to the regulation of extracellular DA dynamics during successive periods of dopaminergic stimulation.

Several convergent mechanisms could account for this neurochemical pattern. Beyond their role in D2R desensitization (Daigle et al., 2014), presynaptic D2 autoreceptors function as the primary feedback regulators of DA synthesis and release (Bello et al., 2011). Furthermore, functional interactions between D2R signaling and DAT activity strongly influence extracellular DA clearance (De Mei et al., 2009). The observed neurochemical disruption could therefore stem from impaired receptor autoinhibition, altered DAT-mediated clearance, a rapid depletion of the releasable vesicular pool driven by the elevated basal tone, or a combination of these processes. While microdialysis provides a measure of net extracellular DA rather than isolating discrete cellular events, our findings suggest that GRK2/3 activity is an additional modulator of DA levels in the DS. Follow-up studies should address the specific mechanisms by which GRK2/3 act in DA regulation.

Interestingly, while the behavioral response to Cmpd101 depended on the baseline novelty-seeking phenotype, its effects on extracellular DA dynamics in the DS did not. This apparent divergence reveals a critical aspect of striatal neurobiology: while local GRK2/3 inhibition changes DA regulation within the DS, the ultimate behavioral consequence is strictly dictated by the individual’s baseline neurobiological state. Highly novelty active animals, characterized by diminished GRK2/D2R availability and elevated baseline DA tone, appear sensitive to further pharmacological inhibition of this pathway. Bridging these molecular, neurochemical, and behavioral domains, our study proposes that the GRK system could be a relevant factor in the definition of a drug addiction susceptible phenotype.

## Acknowledgements

This study was funded by: ANID Fondecyt postdoctorado 3230573 to RIG; Puente UC and Avanza UC 27210068-301-81 to MEA.

