## Supplementary Figures for "Individual differences in novelty-seeking behavior are modulated by GRK2/3 inhibition in rats"

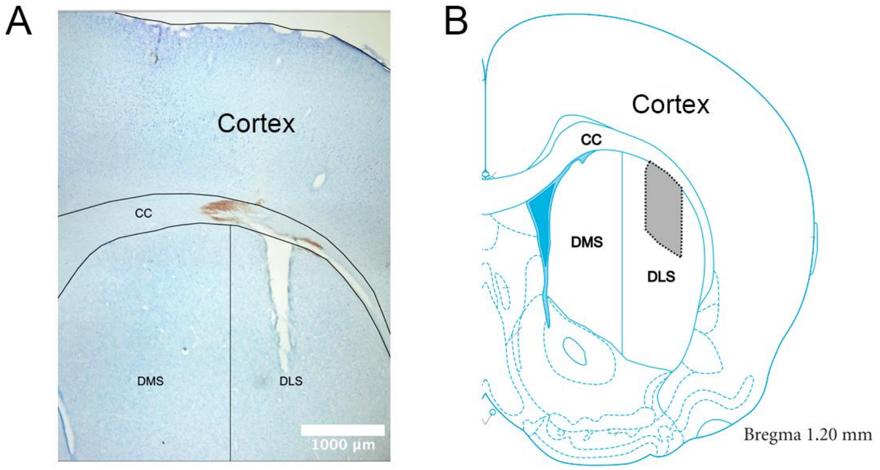


**SUPPLEMENTARY FIGURE 1. Histological verification of the microdialysis probe implantation site in the DLS. A)** Representative photograph of a coronal rat brain section (50 µm) stained with cresyl violet. The vertical mechanical track left by the probe insertion can be observed, confirming its precise anatomical location within the DLS, immediately ventral to the corpus callosum (cc). Scale bar: 1000 µm. **B)** Schematic representation of the coronal section corresponding to the anteroposterior coordinate Bregma 1.20 mm, adapted from the Paxinos and Watson stereotaxic atlas (2009). The gray shaded area delineates the region within the DLS where the active membrane of the probes was positioned. Abbreviations: cc, corpus callosum; DMS, dorsomedial striatum; DLS, dorsolateral striatum.

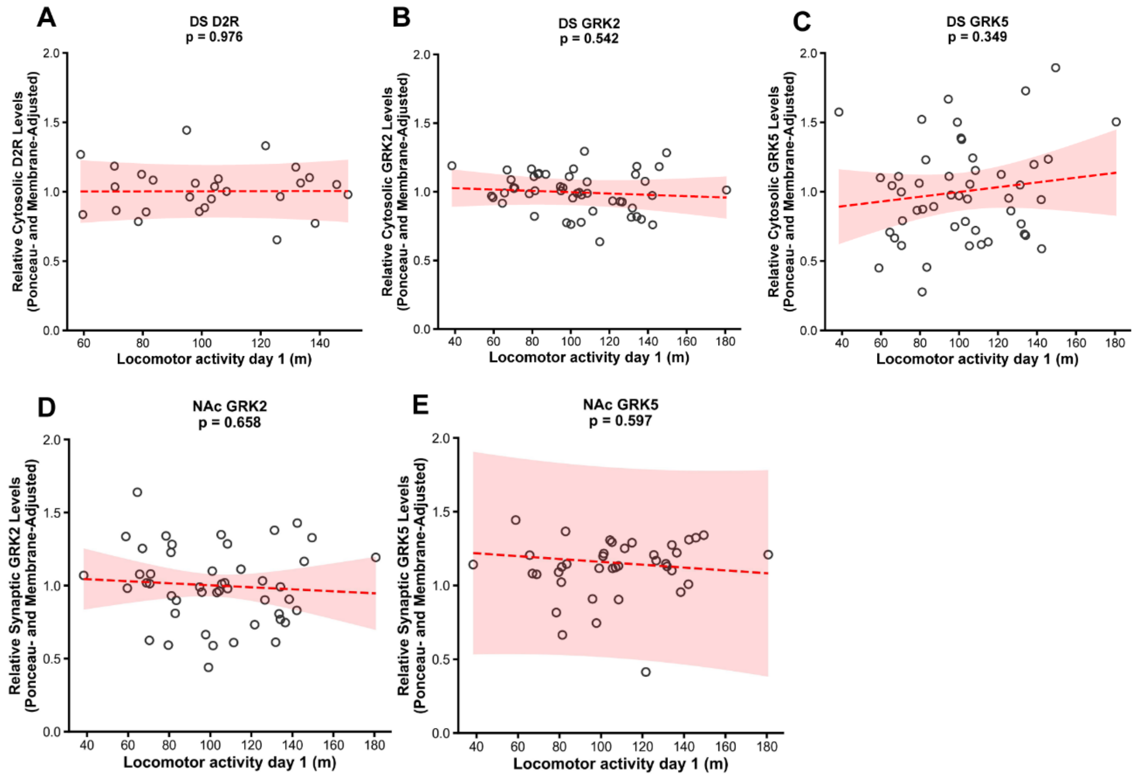


**SUPPLEMENTARY FIGURE 2. Cytosolic D2R and GRKs in the DS, and synaptic GRKs in the nucleus accumbens, do not associate with novelty seeking. A)** Dorsal striatum cytosolic D2R levels do not associate with individual locomotor reactivity (n = 27; p = 0.976). **B)** Dorsal striatum cytosolic GRK2 levels do not associate with individual locomotor reactivity (n = 50; p = 0.542). **C)** Dorsal striatum cytosolic GRK5 levels do not associate with individual locomotor reactivity (n = 49; p = 0.349). **D)** Nucleus accumbens synaptosomal GRK2 levels do not associate with individual locomotor reactivity (n = 47; p = 0.658). **E)** Nucleus accumbens synaptosomal GRK5 levels do not associate with individual locomotor reactivity (n = 40; p = 0.597). Linear mixed models (LMMs) were utilized to account for inter-membrane variability, with Ponceau S signal included as a covariate to control for total protein loading. Dashed red lines indicate non-significant relationships. Shaded areas represent 95% confidence intervals, and each open circle represents one individual animal.

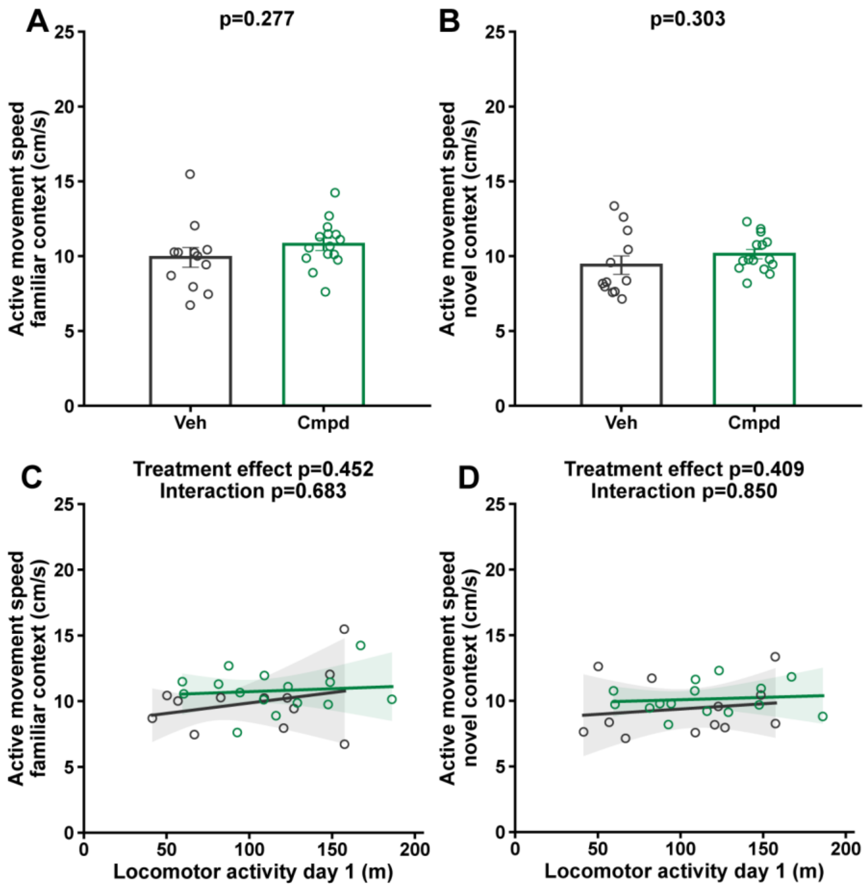


**SUPPLEMENTARY FIGURE 3. Cmpd101 does not alter active movement speed regardless of novelty seeking. A)** Cmpd101 does not affect active movement speed in the familiar context (p = 0.277). **B)** Cmpd101 does not affect active movement speed in the novel context (p = 0.303). Group differences in active movement speed were analyzed using Welch's independent-samples t-test. Bars represent group mean ± SEM. **C)** Active movement speed in the familiar context is independent of locomotor activity on day 1 or Cmpd101 injection (Treatment p = 0.452; Interaction p = 0.683). **D)** Active movement speed in the novel context is independent of locomotor activity on day 1 or Cmpd101 injection (Treatment p = 0.409; Interaction p = 0.850). Baseline-dependent analyses utilized ordinary least-squares (OLS) regression with heteroskedasticity-consistent standard errors (HC3). Black and green lines represent Vehicle and Cmpd101 groups, respectively, and shaded areas represent 95% confidence intervals of the mean prediction. Individual data points are shown across all panels (Vehicle: n = 12; Cmpd101: n = 15). Note: Active movement speed was calculated exclusively during periods of mobility.

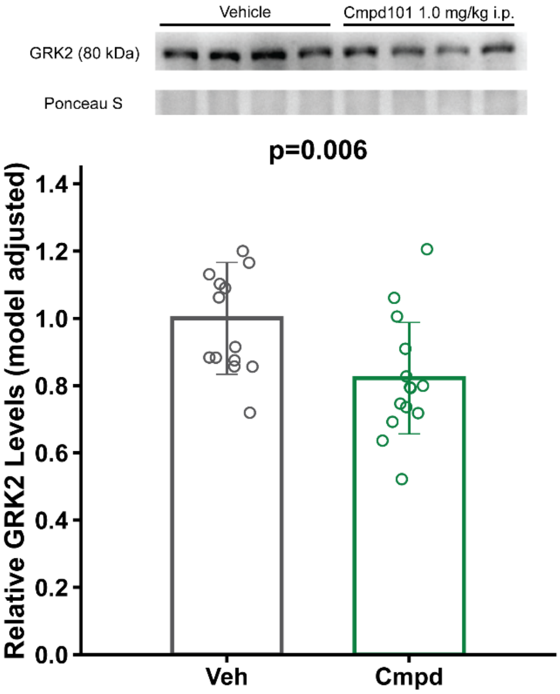


**SUPPLEMENTARY FIGURE 4. Systemic Cmpd101 administration reduces GRK2 protein levels in the DS.** GRK2 protein levels were measured 2 hours after the final vehicle or Cmpd101 1.0 mg/kg i.p administration. Top: Representative GRK2 Western Blots in the DS. Ponceau S staining is shown as the corresponding total protein loading control. Bottom: Relative GRK2 protein levels in the DS. A significant decrease in GRK2 protein levels was observed (Treatment effect p=0.006 using Wald chi-square test). Vehicle: n = 13; Cmpd101: n = 13. Linear mixed models (LMMs) were utilized to account for inter-membrane variability, with Ponceau S signal included as a covariate to control for total protein loading. Bars represent estimated marginal means (EMMs) expressed relative to the reference EMM; error bars represent the corresponding 95% confidence intervals scaled by the reference EMM. Circles represent model-adjusted individual data points expressed on the same relative scale.

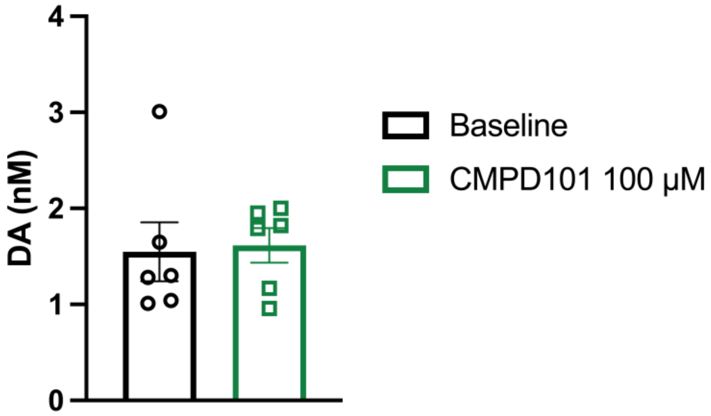


**SUPPLEMENTARY FIGURE 5. Local Cmpd101 (100 μM) perfusion does not alter baseline extracellular dopamine levels in the DS.** Local Cmpd101 perfusion does not affect baseline extracellular DA levels ($p = 0.8366$). Differences in DA concentration were analyzed using a paired t-test. Bars represent mean ± SEM. Black and green borders represent Baseline and Cmpd101 conditions, respectively. Individual data points are shown ($n = 6$). Baseline values represent the average of three consecutive temporal samples, whereas Cmpd101 values represent a single temporal sample collected after at least 30 min of local inhibitor perfusion.
